# Drugs modulating stochastic gene expression affect the erythroid differentiation process

**DOI:** 10.1101/371666

**Authors:** Anissa Guillemin, Ronan Duchesne, Fabien Crauste, Sandrine Gonin-Giraud, Olivier Gandrillon

## Abstract

**Background:** To understand how a metazoan cell makes the decision to differentiate, we assessed the role of stochastic gene expression (SGE) during the erythroid differentiation process. Our hypothesis is that stochastic gene expression has a role in single-cell decision-making. In agreement with this hypothesis, we and others recently showed that SGE significantly increased during differentiation. However, evidence for the causative role of SGE is still lacking. Such demonstration would require being able to experimentally manipulate SGE levels and analyze the resulting impact of these variations on cell differentiation.

**Result:** We identified three drugs that modulate SGE in primary erythroid progenitor cells. Artemisinin and Indomethacin simultaneously decreased SGE and reduced the amount of differentiated cells. Inversely, α-methylene-γ-butyrolactone-3 (MB-3) simultaneously increased the level of SGE and the amount of differentiated cells. We then used a dynamical modelling approach which confirmed that differentiation rates were indeed affected by the drug treatment.

**Conclusion:** Using single-cell analysis and modeling tools, we provide experimental evidence that in a physiologically relevant cellular system, control of SGE can directly modify differentiation, supporting a causal link between the two.

## 1 Introduction

Cell-to-cell variability is intrinsic to all living forms, from prokaryotes [3, 1] to eukaryotes [30]. Such a variability originates from many sources, but arguably stochastic gene expression (SGE) is an important driving force in the generation of cell-to-cell variability among genetically identical cells [9], although additional regulation layers do exist [19]. Classically, SGE is separated into intrinsic and extrinsic sources [13, 26, 2, 25, 33] even if in many cases distinguishing between the two is difficult.

The very existence of SGE led to the concept of a probabilistic mapping between inputs (environment) and outputs (cell decisions) [34]. It is therefore clear that SGE has to be precisely tuned so as to tailor the biological process in which it is involved [11].

Numerous arguments suggest that SGE plays an important role in a wide range of biological processes ranging from bet hedging [36] to the fractional killing of cancer cells [4]. SGE is also involved in decision-making in viruses [39, 38, 40] and in prokaryotes [21, 8], but its role in the differentiation ability of metazoan cells remains an open question. We aim at assessing whether SGE is involved in differentiation or not [20, 18].

If one sees differentiation from the point of view of dynamical systems theory [5], undifferentiated cells are in an equilibrium state at first (self-renewal state). Once differentiation is activated, cells could increase their SGE, exploring a broader region of the state space. Such an exploratory behaviour would increase the probability for cells to attain the space region where they stabilize their gene expression pattern by reaching a new equilibrium state (differentiation state) [18].

We recently described a surge in cell-to-cell variability that accompanies the differentiation of normal primary chicken erythroid progenitors called T2EC [14], that is fully compatible with such a view [28]. Interestingly, these results have been confirmed in various settings, ranging from the differentiation of murine lymphohematopoietic progenitors [23] to the differentiation of murine embryonic stem cells [29, 32].

Nevertheless, a definitive demonstration of a causative role of SGE in differentiation is only starting to appear in the literature [24]. One way of demonstrating this link is the use of drugs that would on one hand modulate SGE and on the other modulate the differentiation process. It has recently been described that such drugs, identified using a large screening approach, were able to modulate the noise affecting the HIV Tat promoter [10]. Recently, drugs that directly inhibit promoter nucleosome remodelling were also shown to provide fine-tuning of SGE [22].

We therefore decided to explore how some of those drugs (Artemisinin and Indomethacin), together with a more general chromatin modifier (MB-3, [26]) could alter differentiation.

Here we show that the three selected drugs significantly modify levels of SGE simultaneously with the level of cell differentiation. We therefore provide the first evidence that in a physiologically relevant cellular system, the modulation of SGE results in a modification of differentiation.

## 2 Results

### 2.1 Drugs affect noise in transcriptomic level

In order to demonstrate a direct link between SGE and differentiation, we modified experimentally SGE in T2EC using three drug treatments: Artemisinin, Indomethacin and MB-3.

Artemisinin and Indomethacin are known to modify SGE of the HIV LTR promoter in human T-lymphocytes [10]. MB-3, a chromatin modifier, is known to modify stochastic gene expression in yeast [26] and in murine ES cells [24]. We first wanted to confirm that these drugs do indeed modify SGE in our cellular system and what are the mechanisms associated with this effect.

We treated T2EC with or without drugs and induced their erythroid differentiation. We then performed single-cell high-throughput RTqPCR on these cells at different time points after differentiation. We assessed a 92 gene panel, relevant for erythroid differentiation study, identical to those previously measured in untreated cells [28]. Single cell transcriptomics data were then analyzed using Shannon entropy as a measure of the heterogeneity among the cells for their gene expression profile [28, 32].

Entropy was affected by all treatments. Under Indomethacin or Artemisinin treatment, entropy significantly decreased after 2 days of erythroid differentiation. This effect was more pronounced with Indomethacin. The opposite effect is observed with MB-3 treatment where entropy was significantly increased after 12h of differentiation for T2EC treated with MB-3 (Figure 1A).

**Figure 1:**
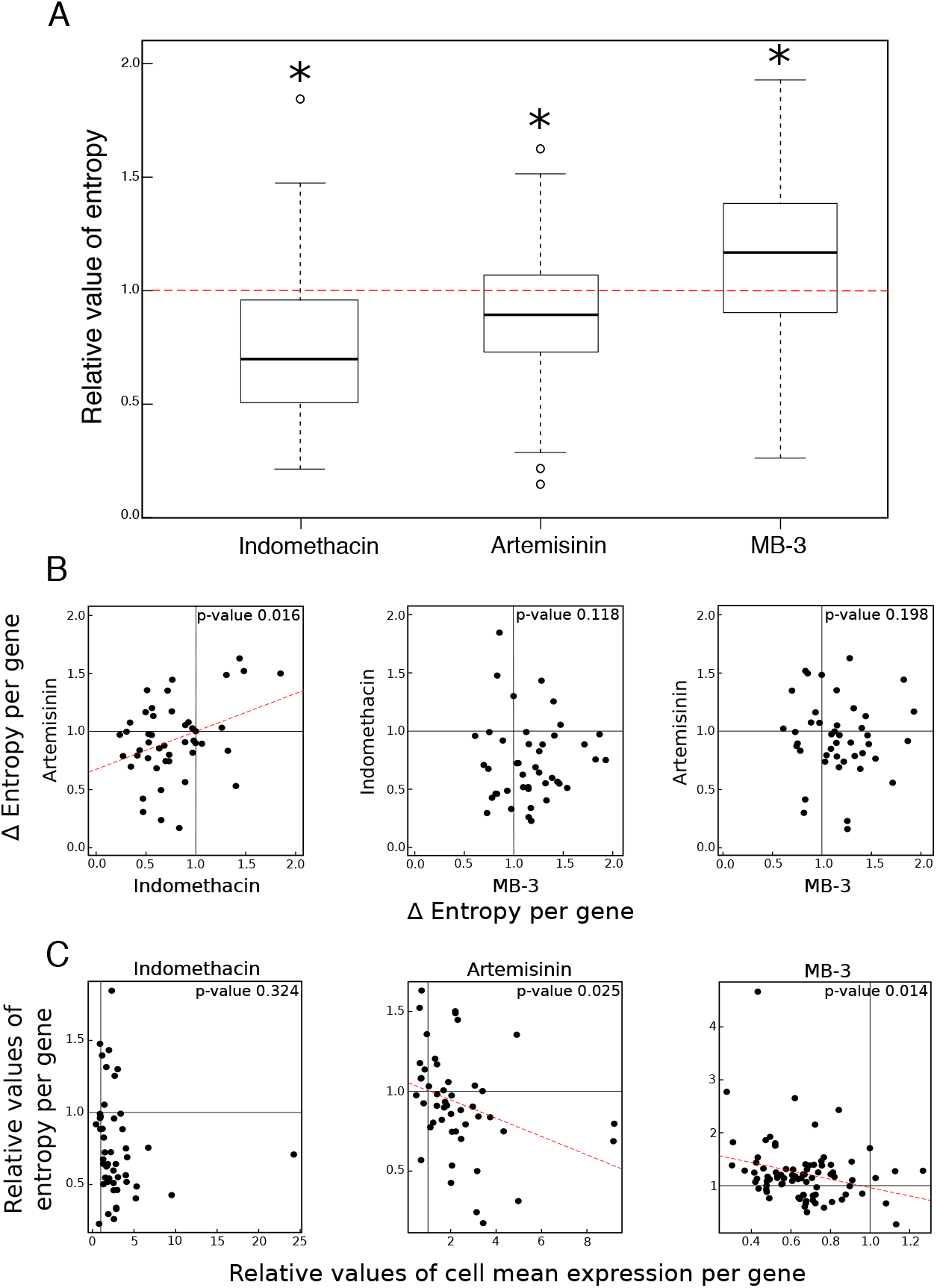
Relative effect of entropy and gene expression average under drug treatment during differentiation. (A) Boxplots representing values of entropy per gene for each treatment relative to control values (red dotted line). Some outliers are not displayed for readability. We assessed the significance of the differences between untreated and treated condition through a Wilcoxon test (tests with a p-value < 0.05 are represented by a star above each boxplot). (B) Correlation plots representing relative values of entropy per gene for each pair of drugs. We assessed the significance of the differences between values for each drug through a Pearson test (p-value < 0.05). When correlation is significant, we displayed the linear regression line for all points (red dotted lines). (C) Correlation plots representing relative values of entropy as a function of relative values of cell mean expression per gene. We assessed the significance of the differences between values for each drug through a Pearson test (p-value < 0.05). When correlation is significant, we displayed the linear regression line for all points (red dotted lines).

We then assessed whether the same genes vary their entropy under the different drug treatments. For this we computed a correlation value between the variations in entropy for each pair of drugs. If the same genes are affected by two drugs, then one would expect their entropy variations to be correlated. We observed a significant correlation only for the genes affected by Indomethacin and Artemisinin treatment. MB-3 treatment seemed to be affecting the variability of a different set of genes (Figure 1B).

The entropy variation could be achieved by modulating the global mean gene expression or the gene expression variance. Thus we finally wanted to test if our drug treatments affected entropy through the modulation of the mean gene expression value. If so, one might expect to see a correlation between the variation of entropy and the mean expression level under drug treatment.

Indeed for two drugs out of three, Artemisinin and MB-3, one observed a significant inverse correlation between mean and entropy (Figure 1C). Nevertheless the effect of Indomethacin on entropy was not related to an effect on mean gene expression.

Here we have found three drugs that modulate SGE in T2EC cells. Indomethacin and Artemisinin decreased it whereas MB-3 increased it. All drugs involve a different set of genes and the effect of drugs was not strongly related to an effect on mean gene expression value. Entropy modulation is therefore the only common characteristic of our three drugs.

We next used these drugs to test their effect on the erythroid differentiation process.

### 2.2 Drugs affect differentiation process

In order to know if drugs modulating SGE also affect the differentiation process, we measured the percentage of differentiated cells in treated and untreated conditions during 96h of erythroid maturation.

A significant modulation in the percentage of differentiated cells was observed for all three drugs (Figure 2).

**Figure 2:**
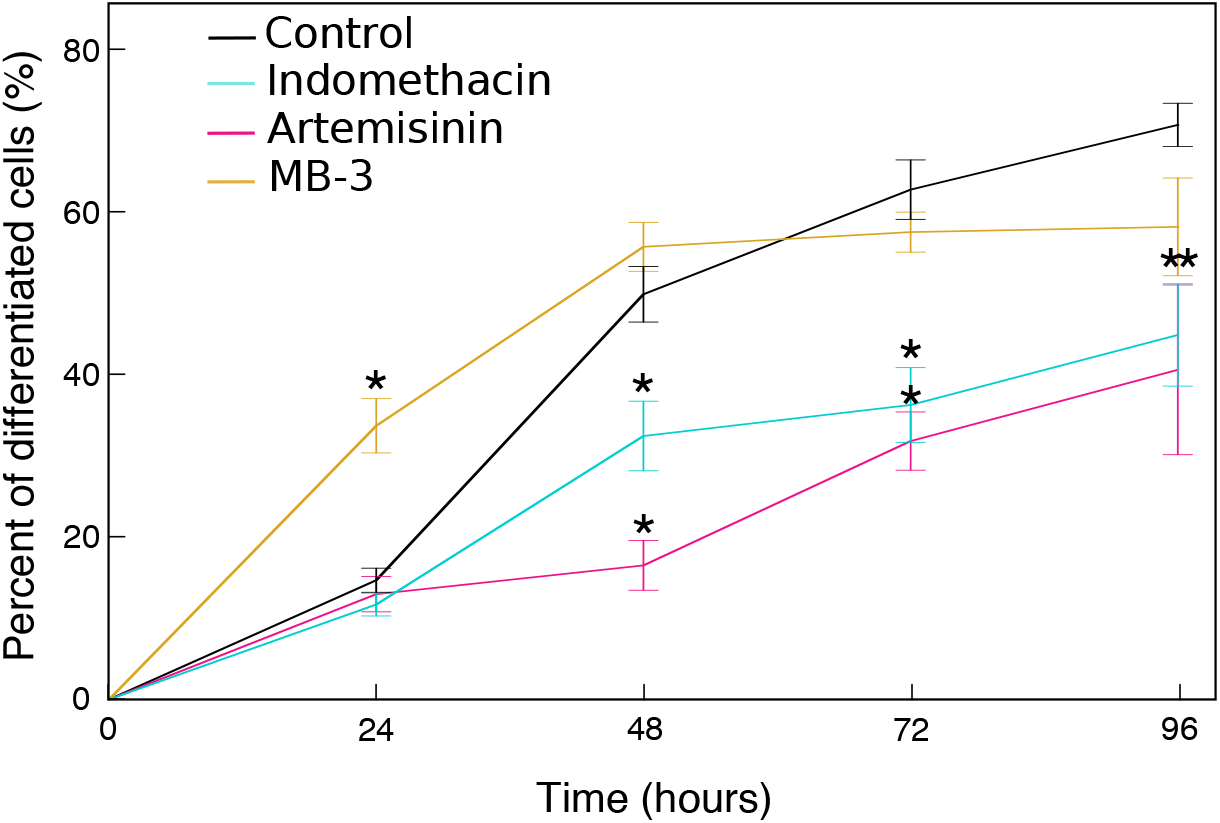
Drugs affected erythroid differentiation. Control conditions were averaged (black line) for readability. Shown is the percentage of differentiated cells for all conditions. Error bars represent the variation between experiments (n=3). We assessed the significance of the differences between each treated condition with their own control condition through a student test (p-value < 0.05).

Indomethacin and Artemisinin decreased the percentage of mature cells from 48h of differentiation onward. MB-3 acted earlier: it significantly increased the percentage of differentiated cells by 24h before returning to somewhat below the control level.

Indomethacin and Artemisinin, two drugs that decreased SGE, reduced the percentage of differentiated cells. Inversely, MB-3 that increased SGE, enhanced the percentage of differentiated cells.

However, at this stage, we cannot conclude that drugs modifying SGE led to a change of the differentiation process itself. Indeed, these effects might have several origins including modification in growth or death rates of our cells. To decipher between these effects, we decided to use a mathematical model describing the dynamics of the *in vitro* erythroid differentiation [12].

### 2.3 Cellular basis of drug effect

Our model describes the dynamics of three cell populations related to three different stages of differentiation. The first one is the self renewing state (S) where differentiation has not started; the third one is the differentiated state (B) where cells have finished differentiating. The second one is the committed state (C), comprising intermediary cells that are committed to differentiation but not yet fully differentiated (Figure 3).

**Figure 3:**
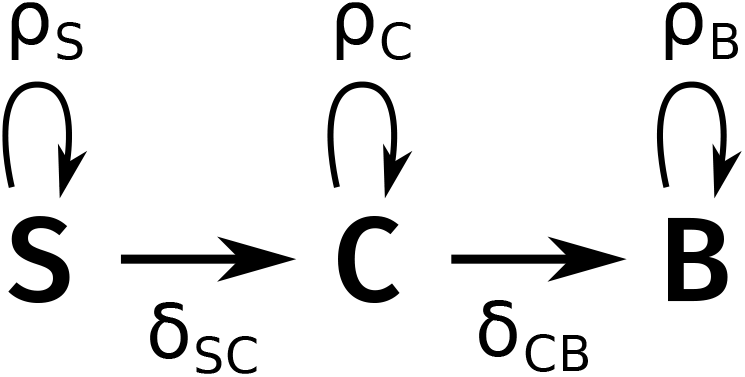
Schematic diagram of the model.

The dynamical model is characterized by a set of five parameters *θ* = (*ρ_S_*, *δ_SC_*, *ρ_C_*, *δ_CB_*, *ρ_B_*):

- *ρ_i_* is the proliferation rate of compartment *i*, involving the balance between cell proliferation and cell death. This value can be either positive (more proliferation than death) or negative (more death than proliferation).
- *δ_ij_* is the differentiation rate of cell type *i* into cell type *j*, which is positive.

Considering that there are no more self-renewing cells after 2 days of T2EC differentiation (Figure S1, [28]), *δ_SC_* is a fixed parameter fully determined by *ρ_S_* [12].

In order to get the best description of the drugs effects with the fewest parameters, we used the same approach as described in [12] and section 4.3.

Using experimental data represented in Figure 2 and following this approach, numerous models are possible for each treatment. Among those, best models are selected by a criterion: Akaike’s weights (Figure S2 & 4.3.3) and reproduced well the cellular kinetics during differentiation observed *in vitro* (Figure S3).

For all of those best models, their parameter values for each treatment are displayed in Figure 4. Under Indomethacin or MB-3 treatment, *ρ_S_* (net growth rate of the immature cells) was not affected in all models and slightly decreased under Artemisinin treatment. Therefore, *δ_SC_* was not affected by the treatments either (data not shown), since its value is entirely determined by the value of *ρ_S_*.

**Figure 4:**
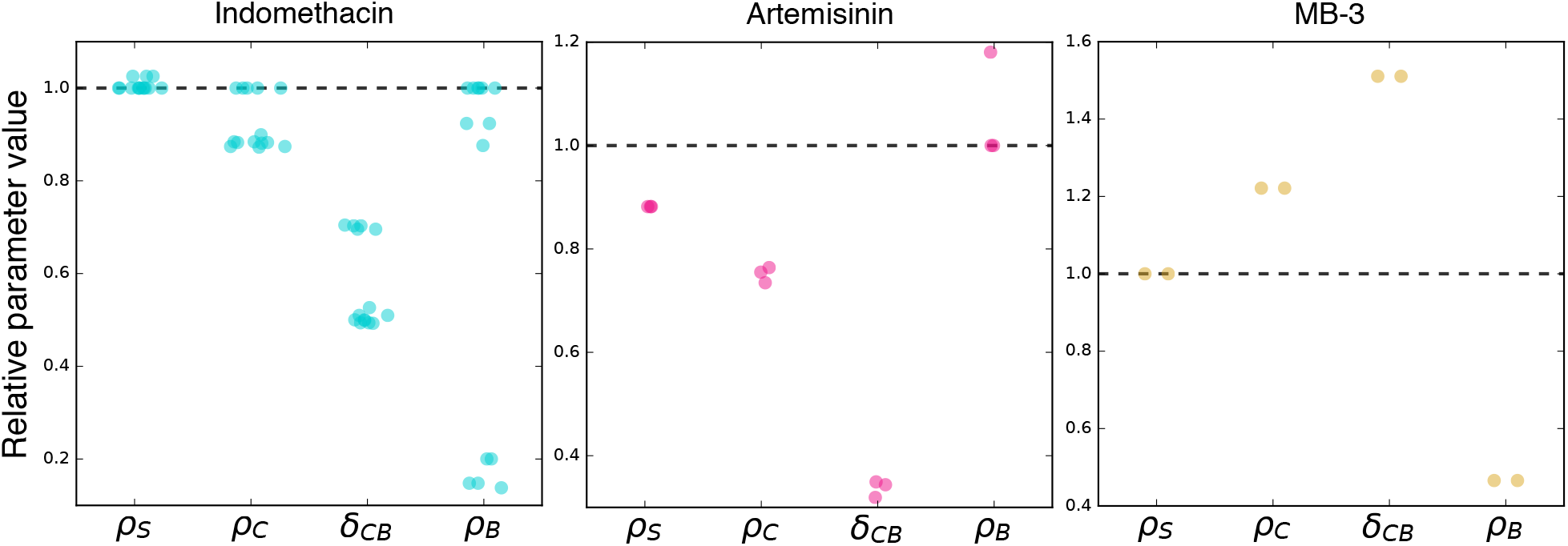
Relative parameter values. For each of the models selected by Akaike’s weights (Figure S2), all the relative parameter values are represented by a dot for a treatment compared to the untreated condition (black dotted line). 14 models were selected for the Indomethacin treatment, 3 for the Artemisinin treatment and 2 for MB-3. The horizontal spacing between the values of each parameter was chosen randomly for readability.

Concerning *ρ_C_*, the net growth rate of the committed compartment, its values were reduced compared to the untreated condition for the majority of models under Indomethacin or Artemisinin treatment, whereas for MB-3 its value increased in both models.

A more variable change between drug effect was observed with parameter *ρ_B_*, which describes the net growth rate of differentiated cells. Under Indomethacin treatment, some of the best models did not show a different value when compared to untreated condition whereas some models displayed a reduced parameter value. Under Artemisinin treatment this value was unchanged for two models among three and increased for the other one. With MB-3 treatment, *ρ_B_* decreased in both models.

Finally, we found that the *δ_CB_* parameter, representing the differentiation rate between committed compartment and mature cell compartment was affected by all three drugs: both Indomethacin and Artemisinin reduced this differentiation rate whereas MB-3 increased it in all best models.

These results demonstrate that all three drugs alter the differentiation process by modifying all dynamical parameters including the differentiation rate between committed and mature cells: it is clear that drugs that reduce SGE decrease differentiation rate and inversely that the drugs increasing SGE accelerate cell differentiation, in line with our initial hypothesis.

## 3 Conclusions & discussion

In this study, we assessed the existence of a direct link between the modulation of stochastic gene expression and a differentiation process. We tested drugs known to modulate SGE in different cellular systems [10, 26]. We showed that these drugs modify the level of SGE in our cells. We therefore tested their effect on the differentiation ability of avian erythropoietic progenitors. We identified which differentiation parameter were affected by drugs using a dynamical model of the *in vitro* erythroid differentiation [12]. We demonstrated that drugs modulating SGE level affected the differentiation process by impacting the differentiation rate between the two last compartments. We therefore demonstrated a direct link between SGE and a differentiation process supporting our starting hypothesis that stochastic gene expression participate positively in cell decision-making to differentiate.

Indomethacin, Artemisinin and MB-3 have clearly different functions. Artemisinin is an anti-malarial drug used against a parasitic infection [15]. Indomethacin is an anti-inflammatory drug that affects the prostaglandin pathway [17]. MB-3 is an inhibitor of GCN5, a histone acetyl transferase (HAT) that activates global gene expression [41]. Even in such a seemingly well-defined case, it should nevertheless be remembered that a very complex relationship may lie between the biochemical action of a drug (HAT inhibition) and its biological effect on SGE [40].

Considering these different functions, it is hard to imagine that all these drugs have in common anything else than their ability to modulate SGE in our cells.

The question then arises of the mechanisms through which these different drugs modulate SGE. We first assessed if these drugs affected the entropy of the same genes. For Indomethacin and Artemisinin, we showed that indeed the entropy of some of the same genes were affected but with a weak correlation. In contrast, MB-3 increased SGE through a different set of genes. This tends to indicate that cell-to-cell variability *per se*, relatively independently of the gene function involved, is participating to the differentiation process (see below).

We then investigated a potential role for variation in the mean gene expression that could explain the SGE level variation.

Modifying SGE level is accompanied by a variation in the mean gene expression level for two drugs out of three. The decrease of mean gene expression under MB-3 treatment has been shown in a diffrent system not to be significant [24]. Also, it has not been reported that Artemisinin affect mean gene expression in any other cellular system. However, the fact that Indomethacin treatment decreased gene-wise entropy clearly without affecting the mean gene-wise expression level reinforces the fact that the modification on the differentiation process is not due to a modification in mean gene expression but only to a non-specific modulation of SGE.

Collectively, these results suggest that neither common genes nor common mechanisms could explain the observed effect of the three drugs simultaneously. This reinforces the fact that modulation of cell-to-cell variability has a strong role in differentiation, independently of gene function or the specific mechanism involved.

This could be explained by adopting a dynamical systems view on the differentiation process, in the wake of Waddington’s proposal [37]. In such a view, we could consider that in the highly dimensional gene expression space, an equilibrium cell state could be compared to a valley in an epigenetic landscape [18]. When we reduce SGE using Indomethacin or Artemisinin, we dig the valley, limiting the ability of cells to escape from a self-renewal equilibrium. Their probability to attain the new equilibrium state is reduced. Inversely, when we increase SGE using MB-3, we flatten the valley and improve the ability of cells to explore a larger dynamical landscape, and increase their probability to attain the new differentiated equilibrium state more quickly. Once cells achieved their journey, they stabilize their new gene expression pattern (the differentiated genetic profile) and return to a basal level of SGE [20, 18, 5]. In such a view, stochastic gene expression favours cells making the decision to differentiate, modifying the structure of the valley in which cells are moving. In a recent perpective, this same process of actively shaping the Waddington Landscape has been described in terms of a Plinko board, whose nail configuration, composition, and patterning can be modified towards forward stochastic design [11]. Similarly to our initial description [28], the variation of cell-to-cell gene expression in other differentiation systems has been recently described [31, 23, 29, 32, 24]. Furthermore, a functional link between transcriptional heterogeneity and cell fate transitions was demonstrated recently through manipulation of the histone acetylation landscape of mouse embryonic stem cells [24]. This is fully backed up by our own data that also demonstrate that the reverse (inhibiting differentiation by reducing SGE) can also be demonstrated.

In recent studies, it has been shown that active modulation of SGE is responsible for a modification of decision-making in viruses [10]. In primary erythroid progenitor cells, we show here that experimentally modifying SGE affects the differentiation process. It could therefore be important to study the potential use of SGE-modifying drugs in differentiation-related diseases such as tumoral cell progression [35], as exemplified by the chronic myeloid leukemia [16, 6, 11], paving the way to a “treatment by noise” of at least some cancer-related diseases.

## 4 Methods

### 4.1 Cell culture and treatment

T2EC were extracted from bone marrow of 19 days-old SPAFAS white leghorn chickens embryos (INRA, Tours, France). These cells were maintained in a medium called LM1. It is composed of *α*-MEM medium supplemented with 10 % Foetal bovine serum (FBS), 1 mm HEPES, 100 nm *β*-mercaptoethanol, 100 U/mL penicillin and streptomycin, 5 ng/mL TGF-*α*, 1 ng/mL TGF-*β* and 1 mm dexamethasone as previously described [14]. T2EC were induced to differentiate by removing the LM1 medium and placing cells into the DM17 medium (*α*-MEM, 10% foetal bovine serum (FBS), 1 mm Hepes, 100 nm *β*-mercaptoethanol, 100 U/mL penicillin and streptomycin, 10 ng/mL insulin and 5% anemic chicken serum (ACS)). Differentiation kinetics were obtained by collecting cells at different times after the induction in differentiation. For Indomethacin and Artemisinin, cells in self-renewing medium are treated at respectively 25 mum and 1 mum 48h before switching into a differentiated medium in order to optimize their effects. For MB-3, cells are treated at 10 mum just after inducing the differentiation. For each drug, a control treatment (0.1% DMSO) was added following the same conditions.

### 4.2 Counting of cell viability and cell differentiation

Cell population growth was evaluated by counting living cells using a Malassez cell and Trypan blue staining (SIGMA). Cell population differentiation was evaluated by counting differentiated cells using a counting cell and Benzidin (SIGMA) staining which stains haemoglobin in blue.

### 4.3 Dynamical model for erythroid differentiation

#### 4.3.1 Calibration, selection and identifiability

This model has been selected among others and has been shown to be a relevant model for the *in vitro* erythroid differentiation process. Moreover, this model is fully identifiable meaning that there is only one parameter set *θ* that corresponds to a dataset [27, 12]. This makes the reasoning on how drugs modify parameter values fully relevant. Details concerning the calibration, the selection and the identifiability analysis of the model are available in [12].

#### 4.3.2 Estimating the parameters under drug treatments

For each parameter of the model, we considered two different cases: one in which the parameter value was unchanged compared to the untreated case (which would not change the number of parameters of the model), and one in which the treatment changed the value (which would introduce a new parameter in the model). Our model has 7 parameters: 5 dynamical parameters presented in 2.3 and 2 error parameters, *b*_1_ and *b*_2_, which quantify the quality of the fit, and the amount of measurement error. These parameters do not influence the dynamics of the model. Only 6 out of these 7 parameters are estimated (*δ_SC_* is set by the value of *ρ_S_*), which defines 2^6^ = 64 models for each drug treatment. We estimated the parameters of these 64 possible models and computed a selection criterion.

#### 4.3.3 Model selection criterion

Estimating the parameters of all possible models under drug treatment needs to be accompanied by the computation of a selection criterion: the model weights based on their corrected Akaike’s Information Criterion 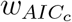 [7]. The Akaike weight of a given model in a given set of models is a measure of the probability that the model is the best one in the set. Thus, selecting the best models of a set of models requires to sort them by their Akaike’s weights. The best models in the set are those whose weights add up to a significance probability (95% in this study) [12].

### 4.4 Single cell high-throughput RTqPCR

Every experiment related to high-throughput microfluidic-based RT-qPCR was performed according to Fluidigm’s protocol (PN 68000088 K1, p.157-172) and recommendations. All the following steps from single-cell isolation to high throughput RTqPCR of each cells are described in [28].

### 4.5 Entropy

We estimated the Shannon entropy of each gene j at each timepoint t as follows: we computed basic histograms of the genes with N = Nc /2 bins, where Nc is fixed for all tests, which provided the probabilities 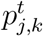 of each class k. Finally, the entropies were defined by

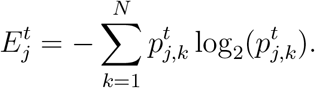

When all cells express the same amount of a given gene, this gene’s entropy will be null. On the contrary, the maximum value of entropy will result from the most variable cell-to-cell gene expression level.

## 5 Acknowledgements

We thank R.D.Dar for his discussions, helpul advices and collaboration. This work was supported by funding from the French agency ANR (SinCity; ANR-17-CE12-0031) and La Ligue Contre le Cancer (Comite de Haute Savoie). We also thank the BioSyL Federation and the LabEx Ecofect (ANR-11-LABX-0048) of the University of Lyon for inspiring scientific events. We thank all members of the SBDM team for fruitful discussions.

